# TTF2 prevents premature rRNA synthesis during mitotic exit

**DOI:** 10.64898/2026.03.22.713493

**Authors:** Catarina Pedro, Laura Tovini, Catarina Peneda, Mia Maesano Krapinec, Raquel A. Oliveira

## Abstract

Mitosis poses major challenges to cellular transcription. As cells enter mitosis, transcription is globally silenced and must be precisely restored upon mitotic exit. These processes are primarily regulated by Cdk1-dependent phosphorylation. In parallel, additional mechanisms, including Transcription Termination Factor 2 (TTF2)-mediated removal of nascent transcripts, reinforce transcriptional shutdown. How these layers of regulation control individual RNA polymerases and influence transcriptional reactivation at mitotic exit remains poorly understood.

Here, we probed how TTF2 differentially controls transcription of distinct RNA classes, using polymerase-specific perturbations and nascent RNA labelling across mitosis. Loss of TTF2 led to accumulation of chromatin-associated transcripts during metaphase, predominantly RNA Polymerase II–derived, consistent with its established role in transcriptional clearance. More unexpectedly, TTF2 depletion caused premature RNA Polymerase I reactivation during anaphase, resulting in unscheduled rRNA synthesis and early recruitment of nucleolar proteins. These findings place TTF2 as a novel regulator of RNA Polymerase I reactivation at mitotic exit. Disruption of this control persists beyond mitosis, resulting in increased nucleolar fragmentation in interphase. Together, these findings reveal TTF2 as a conserved regulator that interfaces with multiple RNA polymerases through functionally distinct modes of control, coordinating both transcriptional shutdown and timely reactivation across mitosis.

## Introduction

Faithful control of gene expression programs across cell division requires not only efficient transcriptional silencing at mitotic entry, but also accurate and timely transcriptional reactivation at mitotic exit. These processes are critical for divisions that maintain cellular identity as well as for those that drive changes in cell fate. Although transcriptional inactivation during mitosis has been recognised for decades ^1^, whether and how the mechanisms that enforce transcriptional shutdown influence the subsequent reactivation of transcription remains poorly understood.

Transcriptional silencing at mitotic entry is driven primarily by the activity of the mitotic kinase Cdk1, which inhibits transcription across multiple RNA polymerase systems. For RNA Polymerase I (Pol I), Cdk1/cyclin B–mediated phosphorylation of one of the SL1 complex subunits, hTAFI110, leads to its inactivation and impairs its interaction with Upstream Binding Factor (UBF). UBF is also suggested to be inactive due to mitotic specific inhibitory phosphorylations, representing an additional layer of regulation of RNA Polymerase I transcription ^2^. In parallel to the regulation of Pol I activity, mitotic entry triggers nucleolar disassembly, in part also mediated by phosphorylation of key nucleolar components. Cdk1-mediated nucleophosmin phosphorylation abolishes its RNA binding activity and thereby modulates its localization and rRNA processing function^3^. Nucleolin is also known to undergo phosphorylation during mitosis, although the functional significance of this modification remains incompletely understood ^4,5^.

RNA Polymerase II (Pol II) is similarly inhibited through Cdk1-dependent phosphorylation of the C-terminal domain (CTD) and components of the general transcription machinery, including TFIID and TFIIH ^6–8^. While these inhibitory modifications efficiently prevent transcription initiation, they are not sufficient to promptly silence ongoing transcription. Accordingly, additional layers of regulation act at mitotic entry to ensure full removal of nascent transcripts from chromatin. Among those, Transcription Termination Factor 2 (TTF2), also known as Lodestar (Lds) in *Drosophila*, has been described as a key regulator of this process^9–12^. TTF2 is a helicase-like ATPase that associates with mitotic chromatin^11,13^ and *in vitro* can displace elongating RNA Polymerases from template DNA ^14–16^. Consistently, depletion of TTF2/Lds leads to retention of elongating Pol II and nascent mRNAs on mitotic chromosomes in *Drosophila* embryos and mammalian cells^9–13^. Although biochemical studies have suggested that TTF2 can also act on stalled Pol I complexes ^16^, its role in regulating rRNA transcription *in vivo* has not been established.In parallel to TTF2 activity, other players have also been proposed to aid mitotic transcriptional silencing, including aurora B-dependent phosphorylation of SAF-A, as well as removal of cohesin from chromosome arms, which triggers the eviction of nascent RNAs and Pol II from chromatin ^17,18^.

Together, these studies define a multilayered network that ensures transcription is efficiently switched off at mitotic entry. By contrast, how transcription is subsequently reactivated remains far less clear. Reactivation could be viewed simply as the reversal of inhibitory phosphorylations and the reassembly of transcription complexes. However, given the extent of chromatin reorganization during mitosis, it is conceivable that transcriptional reactivation follows a regulated and ordered sequence of events, rather than occurring uniformly across transcriptional systems. Accordingly, current models propose that Pol I is reactivated in a multistep manner. Upon mitotic exit, hTAFI110 is dephosphorylated and SL1 rapidly reactivated, driving the initial stages of rRNA transcription in telophase ^19,20^. However, this was shown insufficient for full restoration of maximal Pol I activity, which was proposed to occur later in G1, upon UBF’s activating phosphorylation by G1-specific Cdks/cyclins ^21,22^.

In summary, despite a growing understanding of how Pol II-dependent transcription is silenced during mitosis, how these silencing mechanisms act upon other RNA polymerases, and whether or not they shape the timing, order, and coordination of transcriptional reactivation remains poorly understood. In this study, we investigate TTF2-mediated transcriptional clearance towards different RNA types and uncovered an unanticipated role for TTF2 in restraining premature Pol I reactivation at mitotic exit. Our findings reveal a new layer of regulation that mechanistically links transcriptional inactivation with ordered transcriptional reawakening.

## Results

### TTF2 leads to the accumulation of morphologically different types of nascent RNA across mitotic stages

To investigate the role of TTF2 on transcription dynamics across mitotic stages, we used siRNA depletion in two distinct cell lines, H9 human embryonic stem cells and HeLa cells (Supplementary Fig. 1), coupled to nascent RNAs labelling (5’ ethynil uridine (EU) incorporation) (Fig. 1).

**Fig 1.**
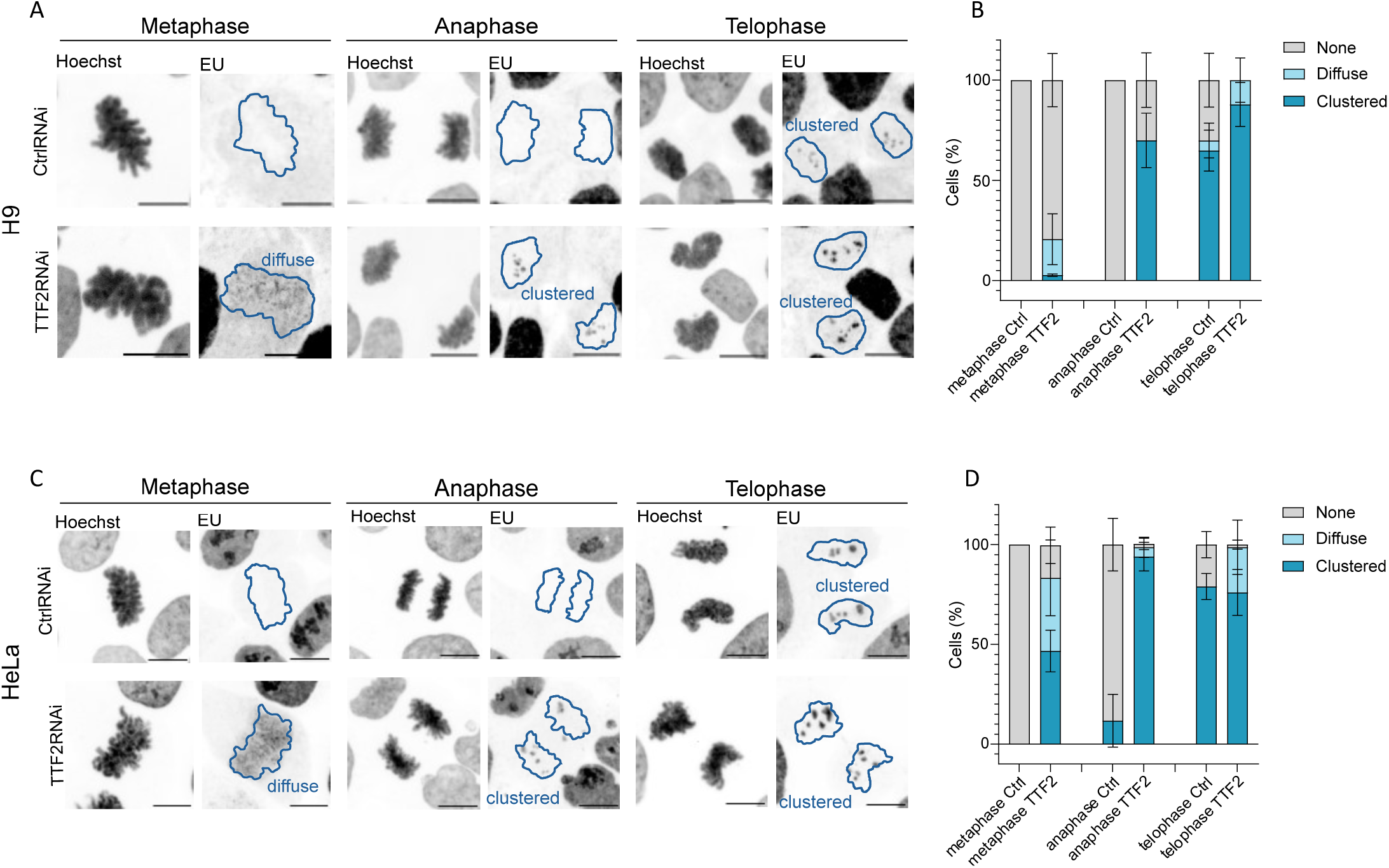
TTF2 knockdown leads to accumulation of nascent RNA transcripts across mitosis in both H9 human embryonic stem cells and HeLa cancer cells. A,C) Representative images of H9 (A) and HeLa cells (C) in Control and TTF2 RNAi conditions with nascent RNA labelling (EU), across the different stages of mitosis. B,D) Percentage of cells with none, diffuse or clustered EU signal across the different stages of mitosis indicated. Data from 3 independent experiments, error bars indicate the mean ± SD. Scale bars are 10 μm.

In both cell lines, we observed that in Control RNAi, EU labelled transcription is virtually absent in metaphase/anaphase cells. In contrast, upon TTF2 depletion, nascent RNAs are present throughout the entire stages of mitosis (Fig. 1). The phenotype was more penetrant in HeLa cells, compared to H9, particularly when considering the total frequency of metaphase cells with EU (82% in HeLa vs 21% H9, Fig. 1D and 1B). Detailed analysis of the distribution of observed EU-signals allowed us to distinguish two distinct phenotypes: a more *diffuse* pattern, characterised by a dispersed signal covering most of the chromatin, and a *clustered* pattern, in which strong signals are confined to discrete sub-chromosomal regions, with little to no signal detected across the remaining chromatin (Fig. 1A and 1C). In both cell types, upon TTF2 depletion, a significant proportion of metaphases exhibit diffuse signals (17% in H9 and 36% in HeLa cells, Fig. 1B and 1D). Clustered signals appear in 46% of metaphases in HeLa cells but are nearly absent in H9 (4%) (Fig. 1C and 1A). Strikingly, anaphase cells show a high percentage of clustered signals in both cell types (69% in H9 and 94% in HeLa).

The presence of a diffuse signal all over metaphase chromatin is compatible with the role of TTF2 in the eviction of Pol II and nascent mRNAs ^9–11^. However, defects in clearance should lead to a constant or decreased EU signal across mitotic progression. Therefore, the high frequency and clustered appearance of EU signals in anaphase is less clear to interpret and suggests a different activity of TTF2 towards specific chromosomal regions. This was particularly striking in H9 cells, where the frequency of anaphase clusters was much higher than the total of metaphases with any detectable EU signal on chromatin (69% vs 20%, Fig. 1B). We therefore focused on exploring the nature of the clustered EU signals during anaphase, using H9 as the prime model system.

### TTF2 depletion triggers an earlier reactivation of ribosomal RNA transcription during mitotic exit

Given the higher frequency of the signals observed during anaphase upon TTF2 depletion, and their clustered appearance, we hypothesised that this signal might correspond to rRNA transcripts, since these are transcribed from repetitive rDNA regions that are clustered in the genome ^23^.

We therefore combined differential inhibition of specific RNA polymerases (triptolide (TRP) for Pol II and BMH-21 for Pol I ^24,25^ with TTF2 depletion. A 2-hour TRP or BMH-21 treatment was sufficient to decrease the amount of nascent RNAs (EU-signal) detected in the nucleoplasm or nucleolar regions, respectively, of interphase cells (Supplementary Fig. 2), confirming the drugs’ efficacy.

We then assessed the consequences of each inhibitor in TTF2-depleted metaphase cells. TRP treatment decreased the amount of metaphase cells that display EU accumulation on the chromatin by around 50%. BMH-21 had a similar effect, albeit to a lower extent (30% reduction) (Fig. 2A and 2B). These findings suggest that nascent transcripts on metaphase chromatin may result from a mixture of mRNA and rRNA transcription events.

**Fig 2.**
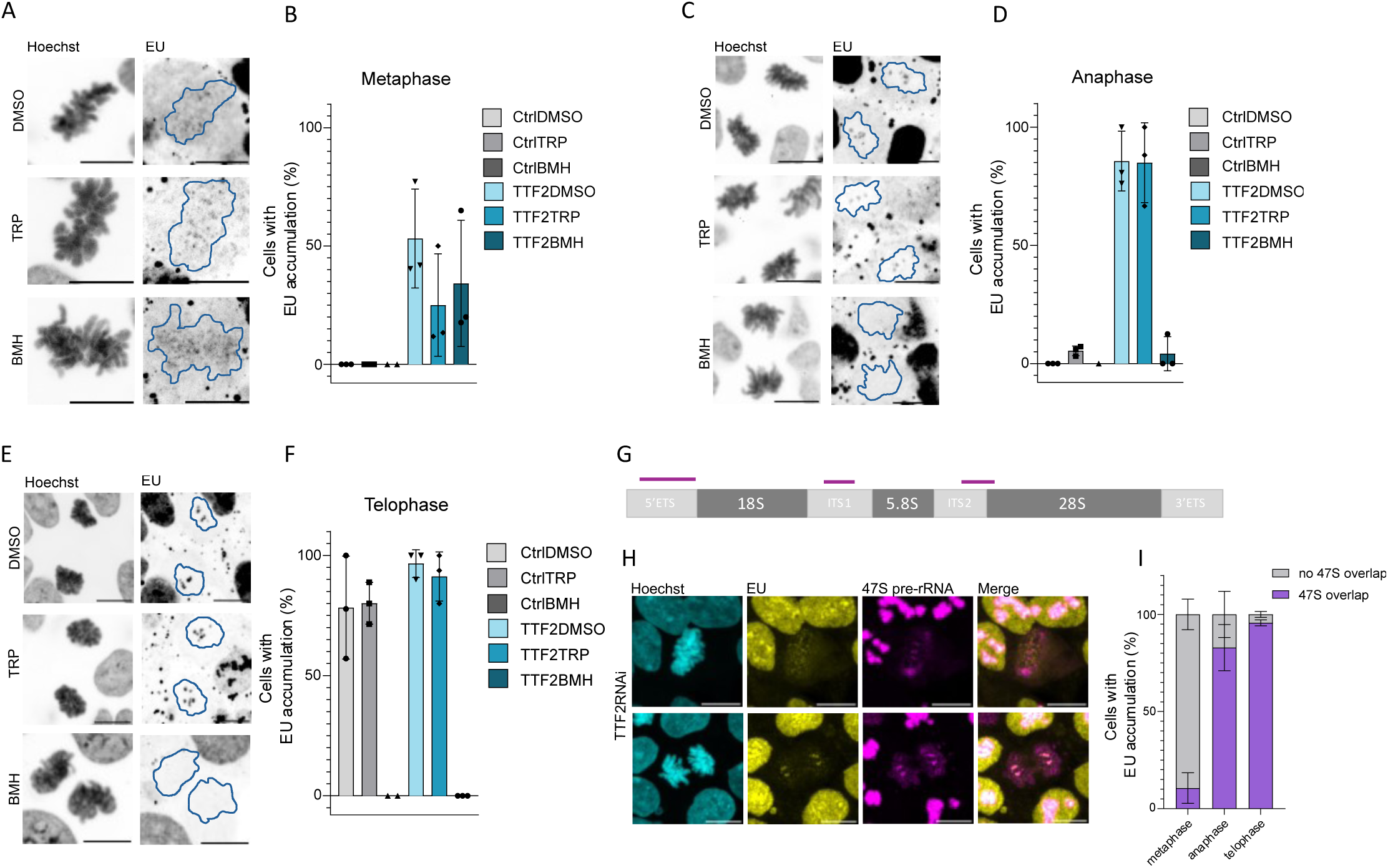
Nascent RNA transcripts in H9 anaphase cells depleted for TTF2 are a consequence of an earlier reactivation of ribosomal RNA transcription. A,C,E) Representative images of nascent RNA labelling in metaphase (A), anaphase (C) and telophase (E) H9 cells in Control and TTF2 RNAi conditions upon DMSO, TRP or BMH treatments. (B,D,F) Percentage of cells displaying nascent RNA labelling (EU). Data from 3 independent experiments, error bars indicate the mean ± SD. G) Schematics of the 47S pre-rRNA gene unit with the sites where the RNA FISH probes bind to (purple line). H) Representative images of TTF2 depleted cells, displaying the overlap between EU and 47S pre-rRNA FISH probes. I) Percentage of cells with nascent RNA labelling with or without overlap between EU and 47S rRNA signal. 2 independent experiments analysed, error bars indicate the mean ± SD. Scale bars are 10 μm.

A similar analysis in TTF2-depleted anaphase and telophase cells produced a much-differentiated response. Whereas TRP treatment had no impact on the frequency of signals detected at these late mitotic stages, BMH-21 fully abolished the presence of nascent transcripts on anaphase and telophase chromatin (Fig. 2C-F). These results indicate that clustered signals result from Pol I-mediated rRNA transcription.

To confirm this possibility, we designed RNA FISH probes that target the 47S pre-rRNA (Fig. 2G). In control interphase cells, this probe shows an overlap with high intensity EU-labelled nascent RNA (inferred as the nucleolus) (Supplementary Fig. 3) confirming its specificity.

We next focused on the analysis of mitotic cells. In TTF2-depleted metaphase cells, we observed very little overlap between nascent RNA (EU-signal) and the RNA FISH probes (∼10% of EU-positive metaphase cells). In contrast, in anaphase cells, this co-localisation occurred in 80% of the analysed cells (Fig. 2H and 2I). These results indicate that upon TTF2 depletion, the RNAs accumulated on metaphase chromatin are poor in Pol I transcripts. In sharp contrast, in anaphase, only Pol I-derived transcripts are detected. Considering the high frequency of nascent rRNAs observed in anaphase cells, despite their nearly absence in metaphase cells (Fig. 2I), we infer that rRNAs detected in anaphase/telophase result from a premature transcriptional reactivation rather than a perdurance of G2 transcripts.

### TTF2 depletion leads to premature transcription of the entire rRNA unit during mitotic exit

So far, our results suggest that TTF2 depletion induces an earlier reactivation of rDNA transcription. However, it is unclear whether this anticipated early onset results in processive elongation across the full transcriptional unit or allows initiation without sustained elongation, leading to transcriptional stalling.

To distinguish between these two possibilities, we designed RNA FISH probes specifically targeting the 5’ External Transcribed Spacer (5’ETS), which marks the 5’ end of the 47S pre-rRNA transcript or the 3’ ETS, corresponding to the 3′ terminus of the transcript (Fig. 3A). To gain further temporal resolution, we focused on anaphase and telophase cells and categorized them as early-, mid-, or late- anaphase (based on the distance between segregating chromatids) and telophase (when chromatin acquired a round shape). In control conditions, the 5’ETS RNA probe is absent in early/mid anaphases and only visible in some late anaphases (14%), and more consistently in telophase cells (84%) (Fig. 3B and 3C). In turn, the 3’ETS signal, indicating transcription of the end of gene, is only visible in about half of telophase cells (Fig. 3B and 3D).

**Fig 3.**
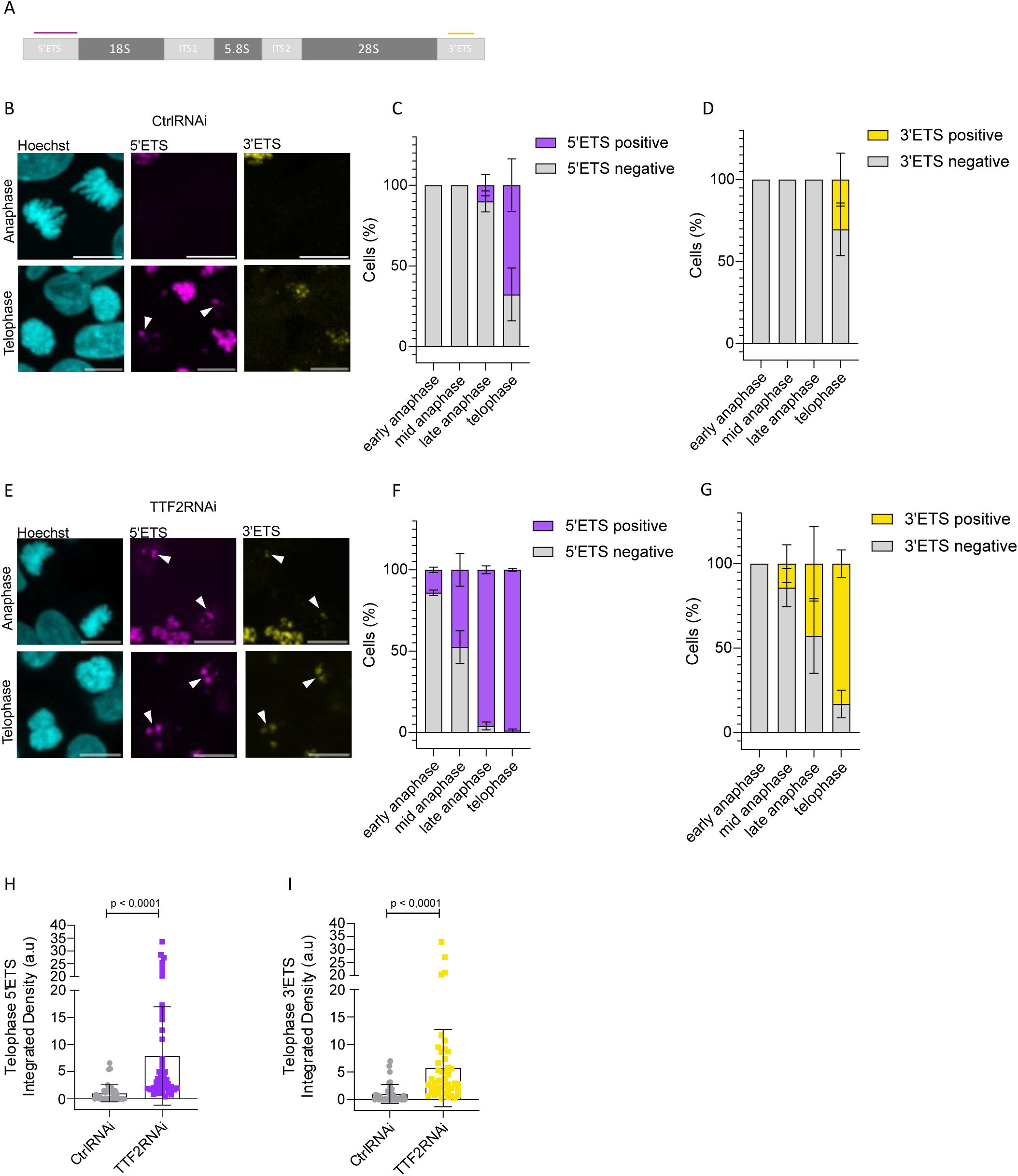
TTF2 depletion in H9 embryonic stem cells leads to an earlier and fully productive reactivation of rDNA transcription. A) Schematics of the 47S pre-rRNA gene unit and the sites where the 5’ETS (purple) and 3’ ETS (yellow) RNA FISH probes bind to. B,E) Representative images of anaphase and telophase H9 cells in Control and TTF2 RNAi conditions, displaying the 5’ETS and 3’ETS. C,D,F,G) Percentage of cells in Control and TTF2 depleted cells that display the 5’ETS signal (C,F), and 3’ETS signal (D,G). Error bars indicate the mean ± SEM from 3 independent experiments. Quantifications of the 5’ETS (H) and 3’ETS (I) signal in Control and TTF2 RNAi telophase cells, where each dots represent one cell. Error bars indicate the mean ± SD of the analysed cells, derived from 3 independent experiments. Statistical analysis was performed using the nonparametric Mann-Whitney test. Scale bars are 10 μm.

When evaluating the time and localization of these same probes in TTF2-depleted cells, we detect expression of the 5’ETS region earlier than in control cells, including some early anaphases (∼12%) (Fig. 3E and F). In agreement with the dynamics of a productively elongating Pol I, the region of the 3’ETS corresponding to the end of the gene can be observed already in mid-anaphase cells (20%) (Fig. 3E and G). Based on these observations, we conclude that not only Pol I starts transcribing earlier upon TTF2 depletion, but also that it can transcribe the entire rDNA gene before cells reach telophase. To estimate the amount of nascent transcripts, in Control and TTF2RNAi conditions, we quantified the signal from each probe. This analysis revealed that in telophase, TTF2-depleted cells display higher levels of both the 5’ETS and 3’ETS probes (Fig. 3H and 3I). We therefore conclude that prematurely reactivated Pol I can transcribe the entire transcriptional unit, leading to higher amounts of full 47S pre-rRNA transcripts produced in telophase cells.

### Premature rRNA transcription is sufficient to trigger partial nucleolar reassembly

Given the known interplay between rDNA transcription and nucleolar assembly ^26,27^, we next sought to evaluate whether premature rRNA transcription, observed upon TTF2 depletion, was sufficient to initiate nucleolar reassembly.

We used two commonly used nucleolar markers, nucleolin and nucleophosmin, to evaluate their co-localisation across the entire EU-labelled region, using only EU-positive cells for this analysis (telophase in controls and anaphase/telophase in TTF2) (Fig. 4).

**Fig 4.**
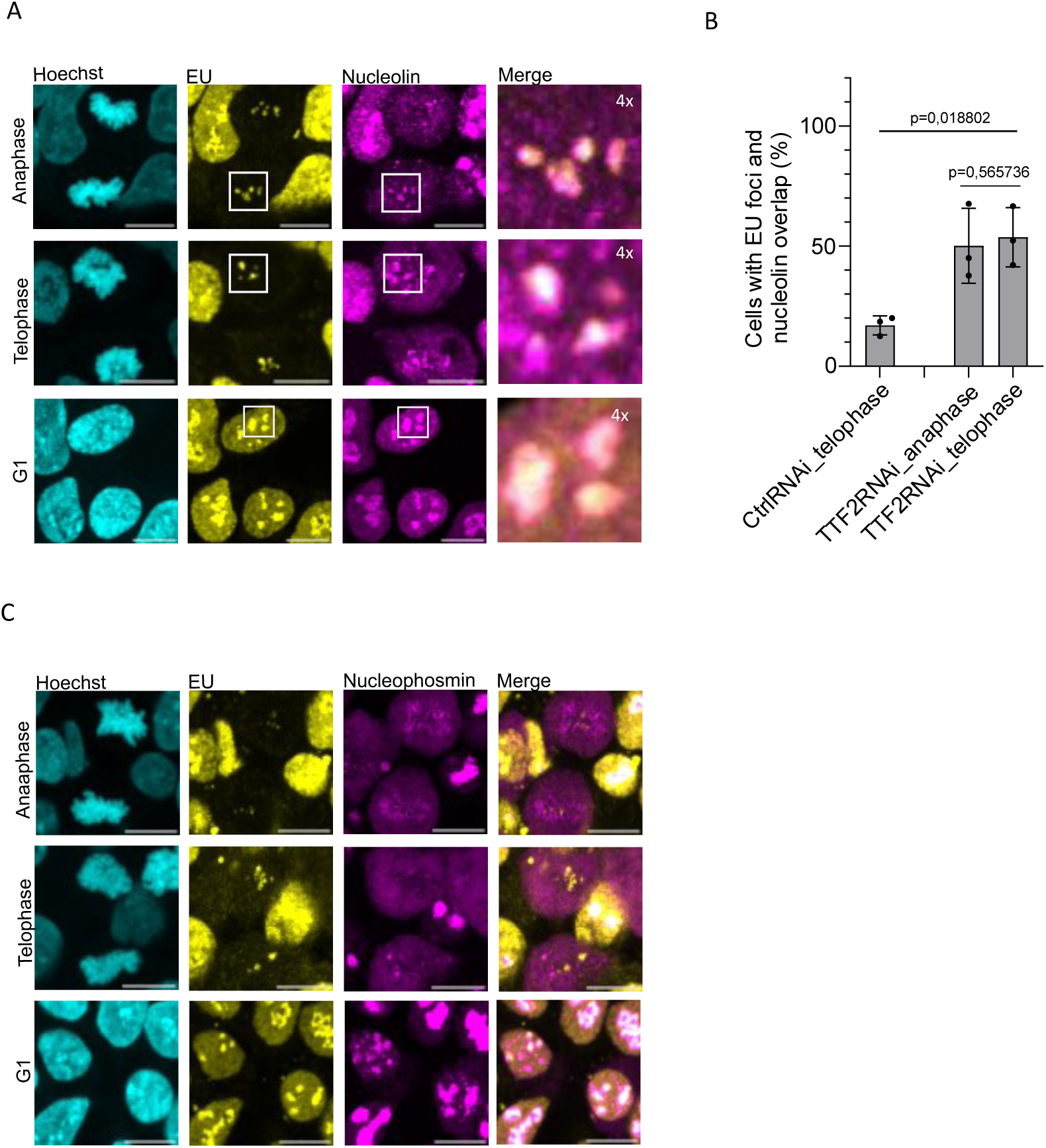
Anticipated rRNA transcription induced by TTF2 depletion is concomitant with earlier recruitment of nucleolin, but not nucleophosmin. A) Representative images of cells with nascent RNA labelling and stained with nucleolin (A) and nucleophosmin (C) in anaphase, telophase and G1 TTF2 depleted cells. B) Percentage of cells that show an overlap between EU and nucleolin, in control telophase cells and TTF2 depleted anaphase and telophase cells. Error bars indicate the mean ± SD from 3 independent experiments. Statistical analysis was performed on cell count, using Fisher’s exact test. Scale bars are 10 μm.

For nucleolin, we observed that in control cells, a full overlap with EU-labelled foci was detected in 17% of telophases. These findings suggest that the recruitment of nucleolar components occurs after the onset of rRNA transcription. For TTF2-depleted cells, these percentages are higher in both anaphase and telophase (50% and 53%, respectively) (Fig. 4A, B). These findings suggest that anticipated rDNA transcription is sufficient to trigger nucleolin recruitment to rRNA transcription sites as early as anaphase.

In contrast, when assessing the localization of nucleophosmin, known to be recruited much later in G1 ^19,28^, we observed no co-localization between EU-labelled regions in anaphase and telophase (Fig. 4C).

These findings suggest that premature transcription of rRNA can initiate the recruitment of some nucleolar components, but not all.

### TTF2 depletion impairs nucleolar integrity in interphase cells

Our findings show that rRNA production, upon TTF2-depletion, is prematurely activated at mitotic exit, with concomitant partial nucleolar assembly. As such, we sought to determine whether this condition affects nucleolar structure beyond mitotic stages.

We first measured the nucleolar area (normalised to the respective nuclear size) of interphase cells, using nucleolin as a marker (Fig. 5A). This analysis revealed no significant differences in nucleolar area between Control and TTF2RNAi (Fig. 5B). However, when determining the number of nucleoli per cell, we observed that TTF2-depleted cells displayed a higher count of individual nucleoli (Fig. 5D,E). Whereas most control cells have 1 - 2 nucleolar bodies upon TTF2-depletion, a high percentage of cells (∼ 60%) display 3 or more nucleoli, indicative of nucleolar fragmentation and/or nucleolar fusion defects.

**Fig 5.**
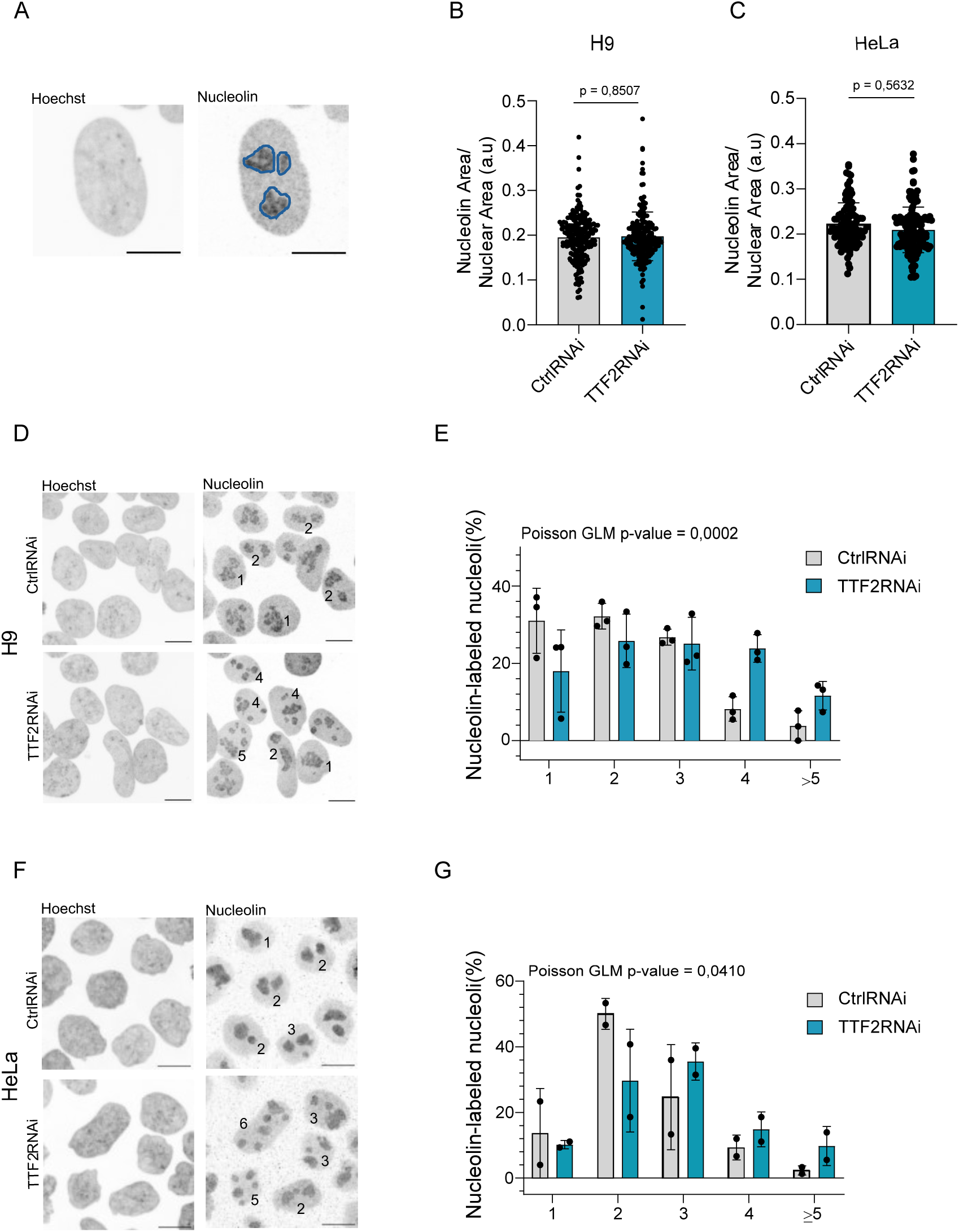
TTF2 depletion in H9 embryonic stem cells and HeLa cancer cells causes an increase in nucleoli numbers. A) Nucleolin staining was used to define nucleolar area and count nucleoli numbers. B,C) Nucleolin area normalised by nuclear area for H9 cells and HeLa cells. Each dot represents one cell, error bars indicate the mean ± SD. D,F) Representative images of H9 (D) and HeLa (F) interphase cells with the indicated number of nucleolin-labelled nucleoli in Control and TTF2 RNAi conditions. E,G) Relative frequency of control or TTF2 depleted cells (H9 and HeLa cells) displaying the indicated number of nucleoli. 3 independent experiments analysed for H9 cells and 2 independent experiments for HeLa. Statistical analysis was performed on cell count, using Poisson generalized linear model (GLM). Error bars indicate the mean ± SD. Scale bars are 10 μm.

To probe whether this phenotype was consistent across different cell types, we performed an identical analysis in HeLa cells. Similarly, no significant differences in nucleolar area between Control and TTF2RNAi was observed (Fig. 5C). When comparing the number of nucleoli displayed by interphase cells, TTF2-depleted HeLa cells also showed an increased number of nucleoli. While control cells mainly display 2 nucleoli, 54% of TTF2-depleted cells have 3 or more nucleoli (Fig. 5F,G), also pointing to nucleolar architectural defects in these cells.

Thus, in the two cell types analysed, TTF2 depletion is linked to abnormal nucleolar organization during interphase.

## Discussion

In this study, we uncover a previously unrecognised regulation of Pol I reactivation, and how it uncouples rRNA transcription from cell cycle progression.

Whereas the role of TTF2 in the clearance of Pol II–dependent transcripts has been previously established ^9–11^, our data reveal a novel role for TTF2 in the regulation of Pol I activity across mitosis, particularly during its reactivation at mitotic exit. We observed evidence for Pol I regulation in both cell types analysed (H9 and HeLa cells), suggesting a conserved role for TTF2 in this process. Evidence for premature reactivation as cells exit mitosis is particularly clear in H9 cells, where Pol I–associated transcripts are ultimately cleared even in the absence of TTF2 (by metaphase), yet reappear prematurely on chromatin during late mitotic stages. In contrast, rRNA transcripts persist on chromatin throughout mitosis in HeLa cells, making it less straightforward to distinguish between delayed clearance and premature reactivation. Nevertheless, the high frequency of Pol I transcripts observed in anaphase, also observed in HeLa cells, strongly supports a conserved role for TTF2 in regulating Pol I activation during mitotic exit in these cells. How TTF2 controls the timing of Pol I activation remains unknown. Clearance of engaged RNA polymerases is consistent with the established *in vitro* activity of TTF2, where it promotes displacement of both Pol I and II from DNA templates ^16^. However, regulation of Pol I may involve additional mechanisms, as Pol I, unlike Pol II, is not fully evicted from chromatin during mitosis ^29^. It is therefore conceivable that TTF2 induces local changes in DNA organisation at rRNA loci, such as altered helical torsion characteristic of SNF2-like helicases ^30^, thereby impairing efficient transcription initiation or elongation without complete displacement of Pol I.

Our findings of premature Pol I–dependent nascent RNA synthesis at anaphase/telophase upon TTF2 depletion indicate that rRNA transcription can be uncoupled from cell cycle progression. This finding challenges prevailing models of Pol I regulation during mitotic exit. Canonical models propose that Pol I activation depends on reversal of Cdk1-mediated inhibitory phosphorylation events; consistent with this, inhibition of Cdk activity by roscovitine in mitotically arrested cells is sufficient to induce rRNA transcription ^31^. More detailed *in vitro* studies have suggested that full Pol I activation requires phosphorylation of UBF by G1 cyclin–dependent mechanisms, a process thought to occur only in G1 ^2,20^. In contrast, our results support a model in which TTF2 provides a critical layer of repression restraining Pol I activity during mitosis. Notably, depletion of TTF2 in H9 cells induces premature Pol I–dependent transcription without obvious changes in the timing of mitotic progression or Cdk inactivation (e.g. no changes in mitotic progression or nuclear envelope reformation (NERF); Supplementary Fig. 4). This indicates that Pol I reactivation can occur in a mitotic context and positions TTF2 as a safeguard preventing untimely rRNA synthesis during mitotic exit.

Our findings also inform the temporal relationship between Pol I reactivation and early nucleolar reassembly. Notably, this process is factor-specific. For nucleolin, we show that its recruitment can be anticipated by premature rRNA transcription, suggesting that nucleolin recruitment is also uncoupled from cell cycle control and is instead regulated by rRNA transcription onset and/or TTF2. In contrast to nucleolin, nucleophosmin is not detected at transcriptional sites during mitotic exit. These observations are in agreement with its late assembly upon mitotic exit ^19,28^.

We further demonstrate that TTF2 depletion is associated with defects in nucleolar integrity. It remains unclear whether the observed nucleolar defects are a direct cause of mistimed rRNA transcriptional reactivation or reflect additional roles of TTF2 in regulating Pol I. However, the fact that TTF2 binds to chromatin exclusively in mitosis ^11^ strongly supports that nucleolar defects are a consequence of mitotic unscheduled rRNA transcription regulation. In either case, the aberrant nucleolar organisation upon loss of TTF2 likely reflects impaired nucleolar function, a phenotype commonly associated with cellular stress and disease states^32^. Given the central role of Pol I transcription in ribosome biogenesis and cell growth, perturbation of this tightly regulated mitotic program may have broader consequences for cells’ proliferative capacity.

An important remaining question is whether similar mechanisms restrain Pol II reactivation during mitotic exit. While our data robustly capture premature Pol I reactivation, the high dynamic range of rRNA synthesis provides a particularly sensitive readout of early transcriptional restart. In contrast, Pol II reactivation involves lower-amplitude, gene-specific transcriptional events that may be more difficult to resolve during the rapid transitions of mitotic exit. It is therefore conceivable that transcriptional clearance mechanisms may also impact Pol II-mediated transcriptional restart in the subsequent cell cycle. Accordingly, previous studies have shown that increasing cohesin stability results in retention of elongating Pol II on mitotic chromatin and in gene expression defects after mitotic exit ^17^. However, because cohesin and its regulators play central roles in 3D genome organisation and transcriptional control throughout interphase^33^, it remains difficult to disentangle direct effects of mitotic transcriptional perturbation from indirect consequences of altered chromatin architecture.

In summary, our work identifies TTF2 as a central regulator of transcriptional dynamics across mitosis, coordinating transcriptional shutdown at mitotic entry and reactivation at mitotic exit. Although governed by a shared regulatory factor, these processes proceed with distinct kinetics and involve different RNA polymerases, underscoring that transcriptional reactivation is not a simple reversal of mitotic silencing. Together, these findings establish mitosis as an actively regulated transcriptional transition and position TTF2 as a key integrator of transcriptional control and nucleolar organisation across cell division. Failures in mitotic transcriptional control may thus have broader implications for how transcriptional programs are improperly re-established during development or underlie nucleolar dysfunction.

## Materials and Methods

### Cell culture and maintenance

H9 and HeLa cell lines were grown at 37°C in 5% CO_2_ incubator. HeLa cells were grown in DMEM High Glucose (L0103, biowest) supplemented with 10% (v/v) Foetal Bovine Serum (FBS) (biowest) and 1% (v/v) Penicillin–Streptomycin (Gibco).

H9 cells were cultured in TeSR E8 Basal Medium supplemented with TeSR-E8 25X supplement (Stem Cell Technologies) on vitronectin (Gibco) coated 6-well plates. For cell maintenance, EDTA based Gentle Cell Dissociation Agent Tryple-Express Enzyme (Gibco) was used and mechanical dissociation applied. For cell seeding and downstream experiment, cells were detached using TrypLE Express and plated in E8 Basal Medium containing Y-27632 (Rock) inhibitor.

### siRNAs transfection for knockdown induction

RNAi was achieved by transfection with 200 ng/ul siRNA (EHU049891 for TTF2 and EHUFLUC as negative control, MISSION® siRNA (Sigma) using Lipofectamine RNAi MAX (Invitrogen) and Opti-MEM™ (Gibco), according to manufacturers’ protocol.

For HeLa cells, 5 × 10^4^ cells were seeded in each of 12-well plates on glass coverslips and knockdown performed for 48h. For H9 cells, 1,2 x 10^5^ cells were seeded in each of 12-well plates and knockdown was performed in 2 consecutive rounds: the first one at the same time as cell seeding, followed by media replacement after 16h (without Y27632) and a second round of transfection after an additional 8h, for a total of 48h.

### Nascent transcription detection and immunofluorescence

Labelling of nascent transcripts was performed by 5-ethynyl uridine (EU) incorporation (1 mM, 30 min in media) followed by detection via ClickiT chemistry (ClickiT® RNA Imaging Kits, Invitrogen C10329) according to manufacturer’s instructions. When combined with immunofluorescence, after Click-iT RNA reaction cells were washed with PBS and incubated with 3% BSA in PBS for 30 min for blocking. Cells were then incubated with primary antibodies for 1h at room temperature (anti-nucleolin, Abcam ab136649, 1:400; anti-nucleophosmin Abcam ab10530, 1:1000), followed by PBS washes and incubation with secondary antibodies (Goat anti-mouse Alexa Fluor 568 Invitrogen, 1:1000). After wash, DNA was stained with Hoechst (1:1000), and coverslips were mounted with Vectashield (Vector H-1000; Vector Laboratories).

### rRNA FISH

To detect rRNA, FISH probes were designed and produced by Molecular Instruments HCR™ (Supplementary Table 1), and hybridization was performed according to manufacturer’s directions. When used in combination with the EU Click-iT protocol, cells were fixed after EU incorporation and permeabilized overnight with 70% EtOH at −20°C. The following day, the Click-iT protocol was performed as above, followed by the RNA FISH staining.

### Western blotting

To confirm the efficient protein knockdown, cells were collected and stored at −80°C until usage or immediately lysed with RIPA buffer (25 mM Tris-HCl pH 7.6, 150 mM NaCl, 1% NP-40, 1% sodium deoxycholate, 0.1% SDS) supplemented with protease inhibitor (Thermo Scientific PIER78429). Samples were quantified with Bradford assay (Bio-Rad), loaded on a 4-12% Mini-PROTEAN® TGX™ Precast Gels (Bio-Rad) and transferred to a nitrocellulose blot membrane. Western blot analysis was performed according to standard protocols using the following antibodies: anti-TTF2 (1:1000, Abcam ab28030); anti-GAPDH (1:2000, Cell Signalling Technology 14C10); rabbit anti-sheep IgG (H&L) (DyLight™ 800 Conjugated, 1:10 000, Rockland 613-445-002); goat anti-rabbit (IRDye® 800CW, 1:10 000, 926-32211).

### Microscopy

Fixed samples were imaged on an Andor Dragonfly Spinning Disk confocal microscope equipped with Andor Sona sCMOS 4 MPix camera using a 1,600 × 1,350 FOV. 60x 1.2 NA water immersion objective was used.

For live-cell imaging cells were seeded in an 8-well Ibidi chamber (Ibidi 80826-90) and transfected as described above, maintaining the ratio of cells, media and transfectant reagents. 48h after transfection, cells were incubated with SiR-DNA (Spyrochrome) in media for 30 min, washed and replaced with normal media, and imaged in a CO_2_ (5%) and T (37° C) controlled chamber. Cells were imaged with a 40X dry objective every 3 min for 8h.

### Image analysis

Images were analysed using FIJI software, in z-stacks or max intensity projections.

For the qualitative assessment of presence or absence of EU, a blind analysis was performed by attributing random file name to the acquired images. To discriminate chromatin bound signals from the chromosome-periphery coating signal, images were analysed by examining each optical section throughout the z-stack.

For the quantitative imaging analysis of the nascent RNA on chromatin (EU) or rRNA (FISH probes), images were analysed after max-intensity z-projection. The signal from the channel of interest was measured using a mask based on DAPI threshold to delineate the chromatin area. Measurements were subtracted by background intensity (measured on an area outside of the cell).

For the qualitative assessment of EU colocalization with rRNA FISH probes (47S, 5′ETS, and 3′ETS), as well as with nucleolin and nucleophosmin, images were examined across z-stacks to distinguish chromatin-associated signals from signals coating the chromosome periphery. EU overlap with nucleolin and nucleophosmin was classified according to their chromatin distribution and spatial coincidence across the chromatin mass.

### Statistical analysis

Statistical analysis was performed mainly using GraphPad Prism 9.0. and details on the statistical tests used are described in the figure legends.

For statistical analysis of the overlap of nucleolin with EU and nucleolar counts, statistical analysis was performed in Jupyter notebook using the scipy.stats Python package.

For nucleolin, cell counts were categorised as “success” (overlap) or “failure” (no overlap) per cell. 2×2 contingency tables were generated for each independent experiment comparing Control and TTF2 RNAi conditions. Fisher’s exact test was applied to each table to assess whether the proportion of successes differed between conditions. To evaluate the overall evidence across experiments, individual p-values were combined using Fisher’s method. The statistical analysis of nucleoli counts was performed using a Poisson generalised linear model (GLM) fitted using the statsmodels Python package.

## Acknowledgements

We thank all members of the Oliveira lab for fruitful discussions. We thank the Imaging Facility of Gulbenkian Institute for Molecular Medicine (GIMM).

