## Supplementary Table 1 for "TTF2 prevents premature rRNA synthesis during mitotic exit"

Table 1 – List of the FISH RNA target sites, number and sequences of the RNA FISH probes (HCR Molecular Instruments) used.

| Target | Probe-binding Sequences |
|---|---|
| 5'ETS | gCTgACACgCTgTCCTCTggCgACCTgTCgCTggAgAggTTgggCCTCCggA |
| | AgCCgCCTgCCggggCCCgCgggCCTgCTgTTCTCTCgCgCgTCCgAgCgTC |
| | gTCTCCgCggggTTgTCCgCCgCCCCTTCCCCggAgTggggggTTggCCggA |
| | CCCCTCgTCTCTCCTCTCCCCgCCCgCCggCggTgCgTgTgggAAggCgTgg |
| 3'ETS | TCCgggCCgggACggggTCCggggAgCgTggTTTgggAgCCgCggAggCggC |
| | CCgCTTCTTCggTTCCCgCCTCCTCCCCgTTCACCgCCggggCggCTCgTCC |
| 47S pre-rRNA | CCTTgCggTgCTCCTggAgCgCTCCgggTTgTCCCTCAggTgCCCgAggCCg |
| | CCgCTCCCgTgCCgAgTCgTgACCggTgCCgACgACCgCgTTTgCgTggCAC |
| | TCCgggggTCggCCTgCggCgCgTgCgggggAggAgACggTTCCgggggACC |
| | ggCgTgggTCgACCTCCgCCTTgCCggTCgCTCgCCCTTTCCCCgggTCggg |
| | ACCTAgCgCgTTCCggCgCggAggTTTAAAgACCCCTTggggggATCgCCCg |
| | CgTggTgTgAAACCTTCCgACCCCTCTCCggAgTCCggTCCCgTTTgCTgTC |
| | CgTCggCCCCggCCgggTggAAggTCCCgTgCCCgTCgTCgTCgTCgTCgTC |
| | gCCCgTCgTgCTgCCCTCTCggggggTTTgCgCgAgCgTCggCTCCgCCTgg |
| | CCCggAgCgggACCgggTCggAggATggACgAgAATCACgAgCgACggTggT |
| | TgCgCCCgAgCgCggCCCggTggTCCCTgCCggACAggCgTTCgTgCgACgT |
| | CggCCgCgTCggggCCTCgCCgCgCTCTACCTTACCTACCTggTTgATCCTg |
| | CggCCgCgACAACCCCACCCCgCTggCTCCgTgCCgTgCgTgTCAggCgTTC |
| | gTCTCCgCggggTTgTCCgCCgCCCCTTCCCCggAgTggggggTTggCCggA |
| | CTTCACgTCCgTTggTggCCCCgCCTgggACCgAACCCggCACCgCCTCgTg |
| | CCgTCCgTCCgTCCgCCgAgCggCCCgTCCCCCTCCgAgACgCgACCTCAgA |
| | gTCCgCgCgTgggTCCTgAgggAgCTCgTCggTgTggggTTCgAggCggTTT |
| | CCCgCCggCCgCgAgAgCCggAgAACTCgggAgggAgACgggggAgAgAgAg |
| | gCgTCgTCggCggTgggggCgTgTTgCgTgCggTgTggTggTgggggAggAg |
| | AgAAgCgAgggTTCCgCCggCCACCgCggTggTggCCgAgTgCggCTCgTCg |
| | AgAgAgAgAAAgAgAAAgAAgggCgTgTCgTTggTgTgCgCgTgTCgTgggg |
