## Supplementary Fig. 1 for "TTF2 prevents premature rRNA synthesis during mitotic exit"

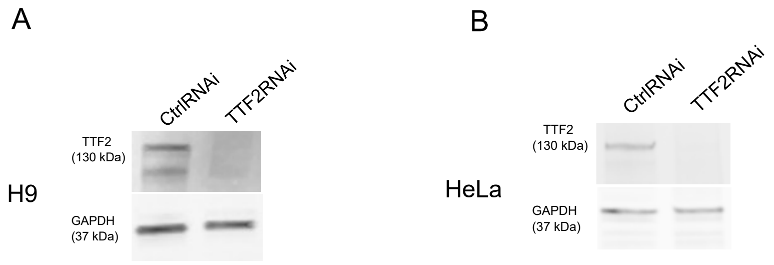


**Supplementary Fig 1. Depletion of TTF2 in H9 human embryonic stem cells and HeLa cancer cells.** A,B) Western blot of Control RNAi and TTF2 RNAi transfected H9 (A) and HeLa cells (B).
