## Supplementary Fig. 2 for "TTF2 prevents premature rRNA synthesis during mitotic exit"

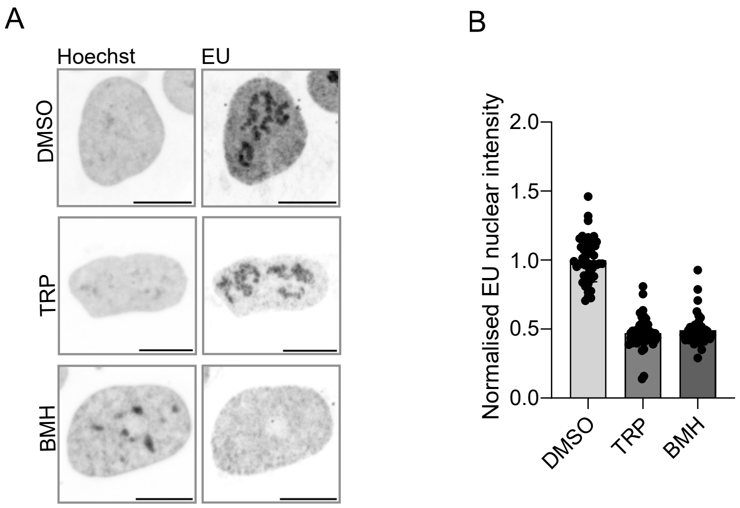


**Supplementary Fig 2. TRP and BMH efficiently inhibit RNA Polymerase II and RNA Polymerase I activity in H9 cells.** A) Representative images of interphase H9 cells in control DMSO, TRP or BMH (2 hour) treatment, showing the nascent RNA labelling (EU) and its decrease on the nucleoplasm and on the nucleolus upon TRP and BMH treatment, respectively. B) Nuclear EU signal intensity normalised to the mean of the control (DMSO). Each dot represents one cell, data from one experiment. Scale bars are 10 μm.
