## Supplementary Fig. 3 for "TTF2 prevents premature rRNA synthesis during mitotic exit"

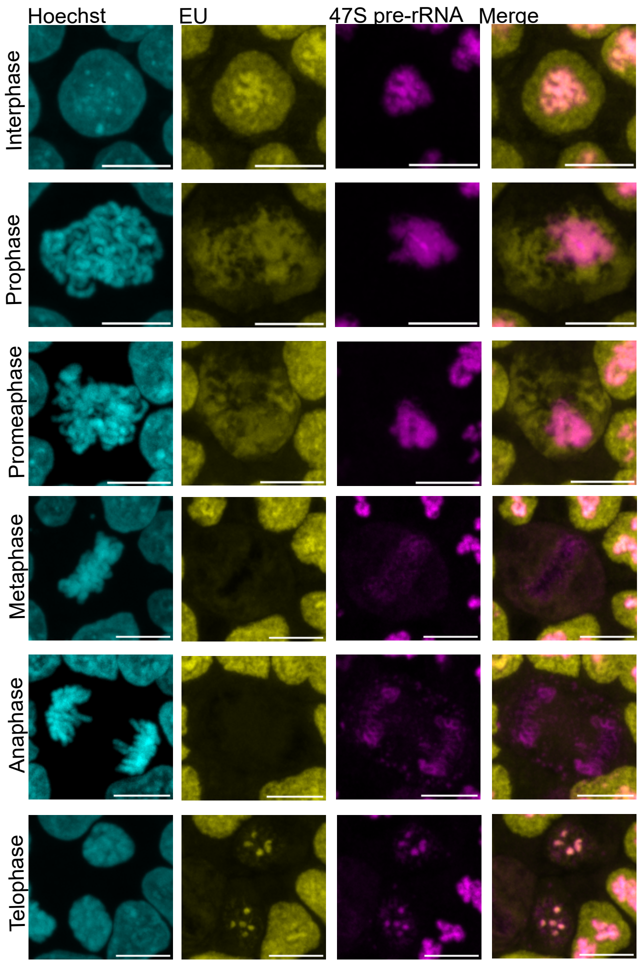


**Supplementary Fig 3. 47S pre-rRNA FISH probes for detection of the ribosomal RNA primary transcript.** Representative images of H9 cells of interphase cells and across the mitotic stages indicated. Scale bar is 10 µm.
