## Supplementary Fig. 4 for "TTF2 prevents premature rRNA synthesis during mitotic exit"

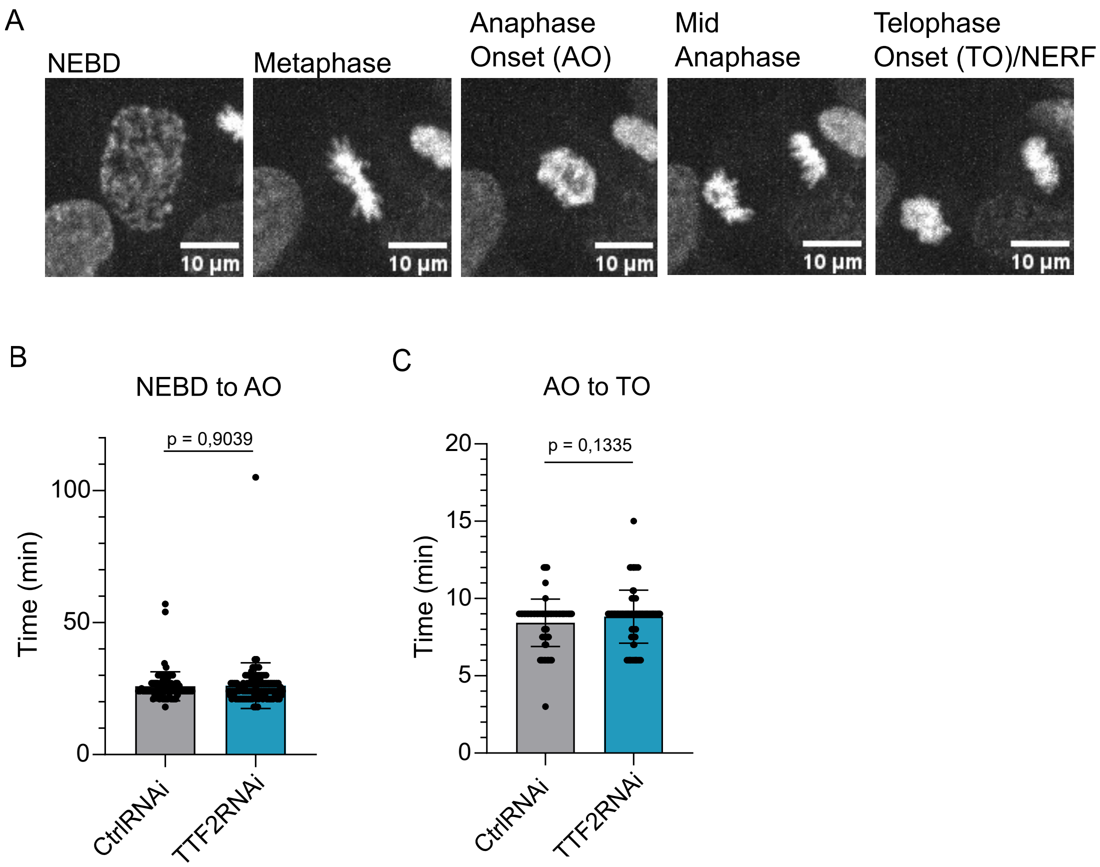


**Supplementary Fig 4. Mitotic timings are not altered upon TTF2 depletion in human embryonic stem cells.** A) Representative mitotic stages of Control cells from live-cell imaging. B,C) Quantification of the time from Nuclear Envelope Breakdown (NEBD) to Anaphase Onset (AO) (B) and from AO to Telophase Onset (TO)/Nuclear Envelope Reformation (NERF) (C) in Control and TTF2 depleted cells. 3 independent experiments analysed. Each dot represents one cell, error bars indicate the mean ± SD . Statistical analysis was performed using the nonparametric Mann-Whitney test.
